# Distribution of soil and clinical *Burkholderia pseudomallei* isolates in Myanmar by using MLST

**DOI:** 10.1101/2024.02.14.580235

**Authors:** Nay Myo Aung, Kyaw Myo Htut, Han Tin Aung, Aye Min Thant, Khine Zaw Oo, Khine Khine Su

## Abstract

*Burkholderia pseudomallei* can be identified as Gram-negative bacillus without spore-forming in an environment such as soil and stagnant water. Human factors and climate changes make the incidence of melioidosis higher. Soil with high moisture and clay-rich is a suitable residence for *B. pseudomallei*. Twenty-one isolates of *B. pseudomallei* were collected and performed by multilocus sequence typing. Among them, eight novel sequence types (STs) of *B. pseudomallei* were found, while the rest show known STs. Interestingly, ST 56 from two patients was identified as the same region, while ST 90 showed local and global distribution. Among six ST 90 *B. pseudomallei*, the three isolates were from soil and patients. Clinical ST 90 isolates were traced in the same region as soil ST 90 isolates. Those isolates may correlate with each other by using multilocus sequence typing (MLST) cost-effectively. However, it still needs to prove its correct relatedness. This study pointed out the reasonable assumption of *Burkholderia pseudomallei* for local and global distribution in Myanmar.

## Introduction

*Burkholderia pseudomallei* can be identified as Gram-negative bacillus without spore-forming in an environment such as soil and stagnant water (1). *B. pseudomallei* can survive in the cells even with the action of many classes of antimicrobial agents, including all macrolides, most first-generation cephalosporins, polymyxins, and aminoglycosides. Fluoroquinolones also are not effective in treatment regimens of melioidosis. *B. pseudomallei*, a causative agent of melioidosis, is a United States of America Centers for Disease Control and Prevention (US CDC) tier 1 select bioterrorism agent. Melioidosis is endemic in Southeast Asia and Northern Australia (3). It is a disease of public health concern and is recognized as a common emerging infectious disease in many tropical countries (4).

Human factors and climate changes make the incidence of melioidosis higher. Soil with high moisture and clay-rich is a suitable residence for *B. pseudomallei* (5). Dissemination of *B*.

*pseudomallei* occasionally occurred in non-endemic regions by implanting infected animals and humans because it can be transported and persist in the environment (7). The geographical distribution of this organism becomes a significant concern in tropical regions. Environmental water study by amplification and genomic sequencing exhibited sporadic cases in India (8, 9). There is challenging to correlate clinical and environmental strains and trace the origin of melioidosis infection. Thailand, a neighboring country of Myanmar, is identified as an endemic area with a high incidence rate (10).

The usage of many molecular typing has been evaluated nowadays. It has been seen in the epidemiology of melioidosis worldwide, consisting of randomly amplified polymorphic deoxyribonucleic acid (DNA) studies, many kinds of ribotyping, and macro-restriction analysis with pulsed-field gel electrophoresis (PFGE). With these many methods, there have been inter-laboratory comparisons, and multilocus sequence typing (MLST) may be chosen to determine the discrimination and search for a clonal relationship. Seven housekeeping genes (seven loci) can be checked in the MLST library from the Pubmed website. MLST screening strategy and datasets have also been illustrated for most highly infectious pathogens, including *Neisseria meningitides, Streptococcus pneumoniae*, and *Staphylococcus aureus* (12). MLST technique becomes a strong discrimination typing of species and is valuable in an epidemiological study cost-effectively. *B. pseudomallei* was first found in a 40-year-old drug-addicted patient in Myanmar by the British pathologist Whitmore AK and his assistant Krishnaswami CS who proved and identified as *B. mallei*-like species in the guinea pig by injection of the isolate from post-mortem infected patient in 1912. They discovered the history and habits of the patient who was released from jail recently. Nevertheless, they thought there was little connection between the transmission mode and infection (13). Surveillance of similar isolates was done, and it was found that 38 cases presented as *B. mallei*-like organisms in the liver, lung, and spleen (14). For this reason, Myanmar is likely an endemic region. However, it is still necessary to perform an epidemiological study on whether it originated from soil or other environmental sources and might be the first origin in Myanmar. This study evaluated the clonal association between soil and clinical isolates by multiple molecular typing methods and genetic analyses in Myanmar.

## Methods

Clinical samples (pus, urine, and sputum or blood samples for clinical isolates) were collected from collaborating centers in different regions of Myanmar. All selected samples were subcultured with Ashdown media at 40°C in the air for 4 days and checked the colonies on each day (15). Following this, isolates that can grow and show characteristic colonies on Ashdown media were tested with a latex agglutination test with safety precautions. After positive colonies were described as *B. pseudomallei* showing agglutination and kept in trypticase soy broth with 20% glycerol for molecular identification (16), soil samples were gathered during September 2018 (the wet season) from a region of disused land and cultivated land in Yangon. Soil culture for *B. pseudomallei* was done according to the protocol (17)

Genomic DNA was obtained from all strains resulting from Vitek2, API 20NE, or latex agglutination tests. Whole-cell suspensions of isolates were utilized for PCR. They will be cultured by plating one loop (1 μl) of stock cell suspension on Trypticase soy agar with 5% defibrinated sheep blood (Himedia). After they had incubated aerobically for 1 to 2 days at 37°C, a single colony was suspended in 200 μl of distilled water in a 1.5-ml Eppendorf tube. DNA of each isolate was extracted with a commercial kit according to manufacturer instructions (Qiagen) and kept at -20°C before amplification. PCR assays were conducted using the target of Type III secretion systems (T3SSs) for species confirmation. PCR optimization and primer design were followed by published articles (18-20). MLST was performed to determine the molecular epidemiology of melioidosis. The following pairs of primers were amplified with the published housekeeping gene fragments (*ace, gltB, gmhD, lepA, lipA, narK, ndh*) (12, 21). The primer pairs designed for PCR amplification were downloaded from the *B. pseudomallei* MLST scheme described in PubMLST (https://pubmlst.org/bpseudomallei/).

PCRs (reaction volumes, 50 μl) were performed with an initial denaturation at 95°C for 4 min, continued by 30 cycles of 95°C for 30 s, 60°C for 30 s, and 72°C for 60 s. The samples were then kept at 72°C for a further 10 min, cooled to 4°C, and stored at −20°C. Amplicons were purified, and Sanger sequencing was performed by First Base Sequencing Service (12).

The sequences were first made alignment and trimmed by Mega 7. Basic sequence assessments such as the number of different alleles, the number of variable positions per allele, and the number and frequency of single nucleotide polymorphism (SNPs) in each locus were assessed by DNAsp V5.1 software (22). Each isolate was analyzed by a string of seven integers (the allelic profile), which correspond to the allele numbers at the seven loci, in the order *ace-gltB-gmhD-lepA-lipA-narK-ndh*. Next, each unique allelic profile was considered a clone and was assigned a sequence type (ST), which also gave a convenient descriptor for the clone. An MLST database containing the sequences of all alleles, the allelic profiles, and information about the *B. pseudomallei* isolates, together with analysis tools, was recorded at Imperial College (London, United Kingdom) and can be examined on the *B. pseudomallei* pages of the MLST website (www.mlst.net). The resulting sequences at the seven loci were concatenated in the order of loci used to determine the allelic profile.

### Ethical statement

Ethical approval was given by Defence Services Medical Academy, Myanmar, Ethical approval number 11/Ethics 2018 for project entitled: Epidemiology of melioidosis and molecular characterization of *Burkholderia pseudomallei* in Myanmar. Ethical approval number 546/2562(EC1) was also given by Siriraj Institutional Review Board, Thailand, for same project entitled: Epidemiology of melioidosis and molecular characterization of *Burkholderia pseudomallei* in Myanmar.

## Result

This study screened 16 clinical and 5 soil *B. pseudomallei* isolates out of 51 *Burkholderia* species in 2018 (Figure 1). Seven housekeeping genes of those isolates were amplified using optimized annealing temperature and described primer in Pudmlst (Table 1) (Figure 2).

**Figure 1.**
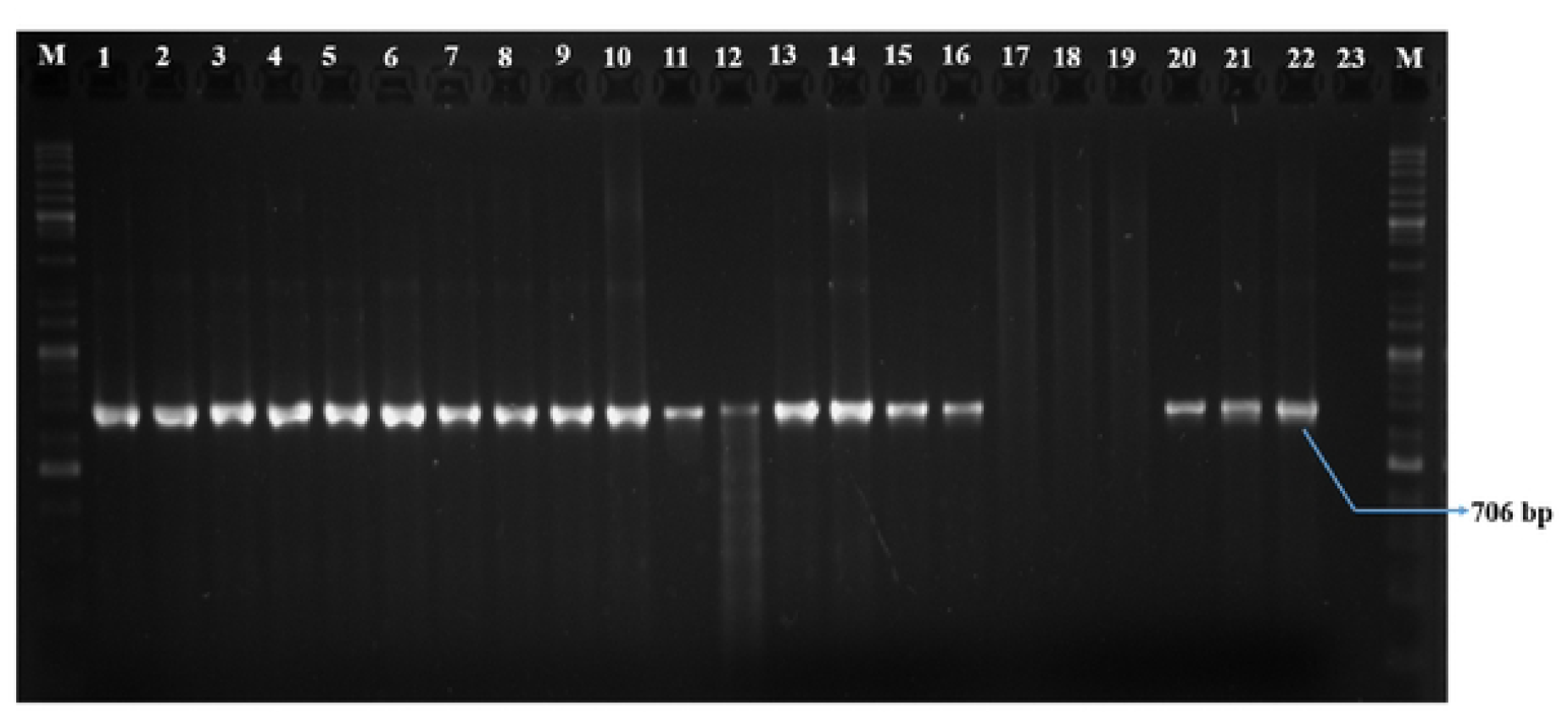
Identification of clinical *B. pseudomallei* by using *ttss* PCR. **(M:** 100 bp and above ladder, lane I: MMBPOOI, lane 2: MMBP002, lane 3: MMBP003, lane 4: MMBP004, lane 5: MMBP005, lane 6: MMBP006, lane 7: MMBP007, lane 8: MMBP008, lane 9: MMBP009, lane 10: MMBPOIO, lane 11: MMBPOl l, lane 12: MMBP012, lane 13: MMBP013, lane 14: MMBP014, lane 15: MMBP015, lane 16: MMBP016, lane 17: MMB006, lane 18: MMB005, lane 19: MMB003, lane 20: MMBOOI, lane 21: MMB002, lane 22: MMB004, lane 23: Negative control, **M:** 100 bp and above ladder)

**Figure 2.**
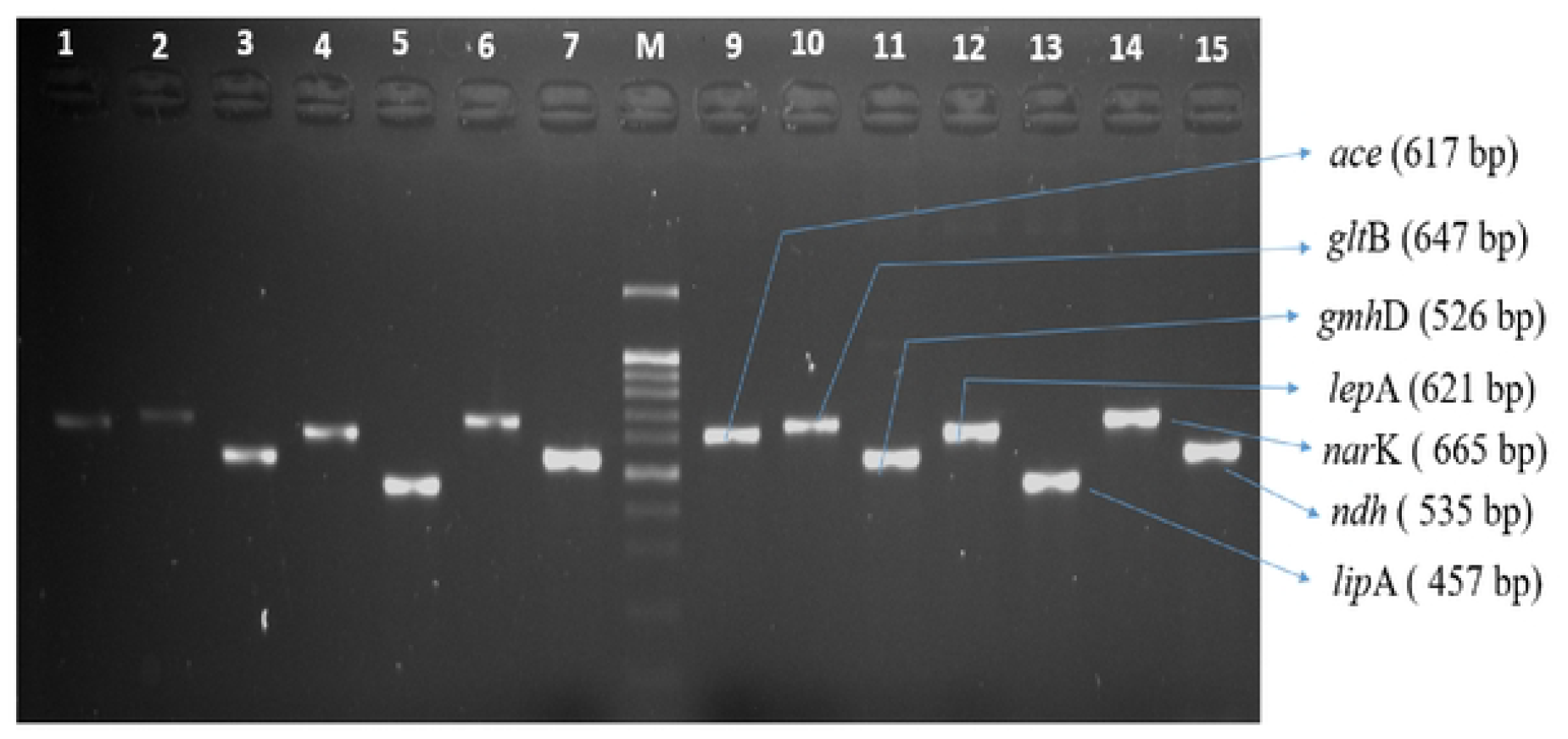
Amplification of seven housekeeping genes of MMB002 and MMB004. Lane I: *ace* (617 bp), Lane 2: *gltB (641* bp), Lane 3: *gmhD* (526 bp), Lane 4: *lepA* (621 bp), Lane 5: *lipA* (535 bp), Lane 6: *narK* (665 bp), Lane 7: *ndh* (535 bp), Lane M: 100 bp above ladder, Lane 9: *ace (611* bp), Lane 10:*gltB* (647 bp), Lane 11: *gmhD* (526 bp), Lane 12:*lepA* (621 bp), Lane 13: *lipA* (535 bp), Lane 14: *narK* (665 bp), Lane 15: *ndh* (535 bp)

Among 21 isolates, allele variations per locus were recorded and ranged from 2 to 5. The level of sequence locus diversity for each of 7 housekeeping genes was approximately 6% (Table 2). Out of 21 isolates, ST 90 (n=6, 28.57%) was found as common ST from 3 clinical and soil isolates, respectively (Table 3). The remaining isolates were resulted as previously published and uploaded sequence types ST300 (n=1, 4.76%), ST 56 (n=2, 9.52%), ST 354 (n=2, 9.52%), ST 416 (n=1, 4.76%), which were isolated from clinical samples, whereas soil isolate showed ST 42 (n=1, 4.76%). The rest 8 isolates were identified in novel STs, representing ST 1722, ST 1723, ST 1724, ST 1725, ST 1727, ST 1728, and ST 1729 from clinical samples and ST 1726 from soil samples (Table 3).

### Correlation of STs in Myanmar with global STs

After PHYLOViZ analysis, studied STs were not found within the same group, showing different clonal relations among regional STs. In addition, Myanmar STs 90 were found not only in patients but also in the environment. In contrast, soil STs 90 (n=3) were detected only in Yangon (the former capital of Myanmar). However, clinical ST 90 (n=1) was distributed in Yangon, Naypyitaw, and InnMa townships around lower Myanmar, respectively (Figure 3). The weather was not noted during the study period, but most had similar conditions in those regions.

**Figure 3.**
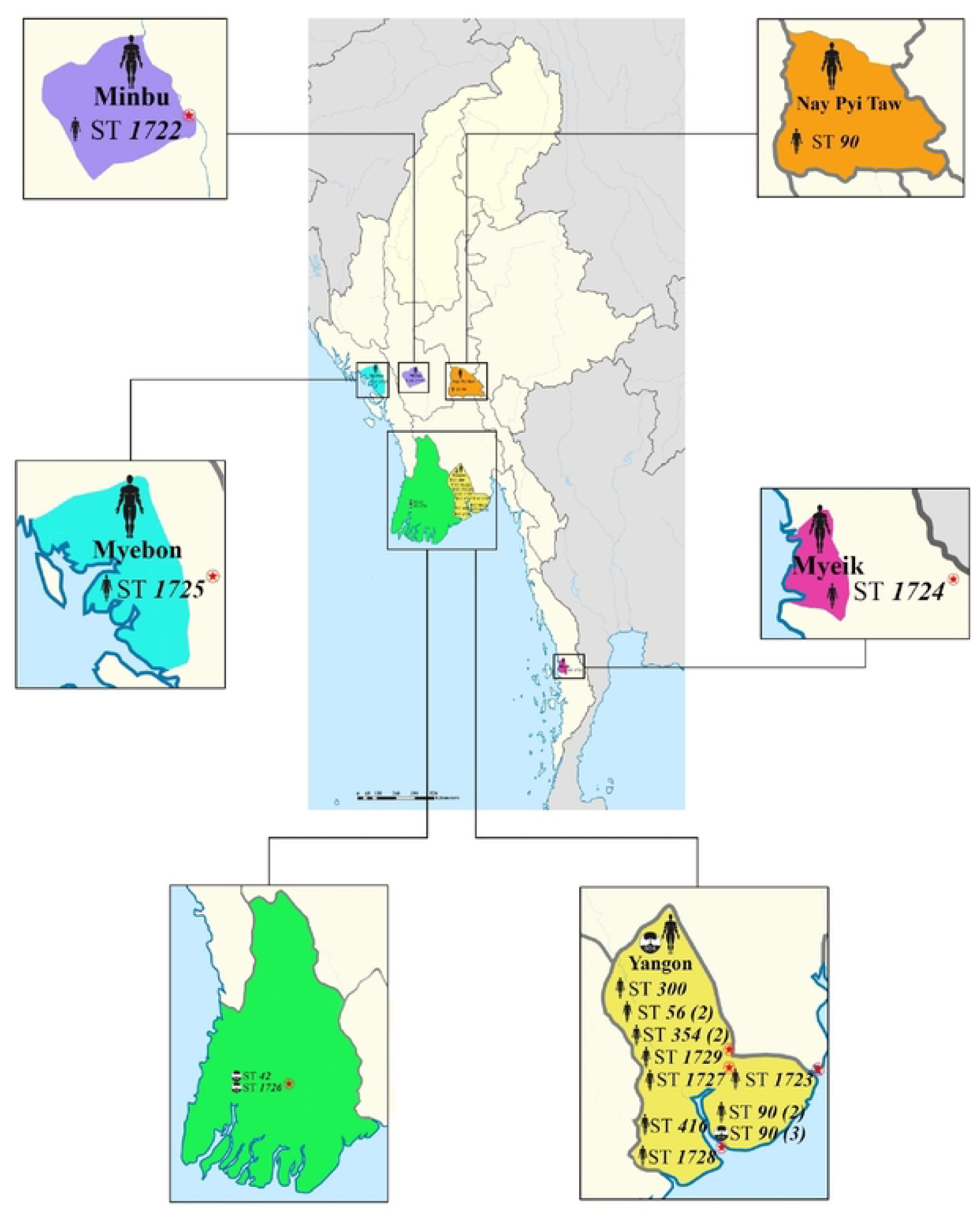
Distribution of Clinical and Soil *Burkholderia pseudomallei* STs with each colors representing different regions around Myanmar. (): number of same ST, Asterisk: novel ST, 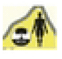: Soli and Human

There was a relationship between STs in this study, and a global collection of 5905 isolates in the *B. pseudomallei* MLST isolates database (as of 4^th^ October 2019) by using PHYLOViZ analysis. All STs in this study were classified into 6 groups, showing the same group with STs from other countries. Group II (ST 300), VI (ST 42), and I (ST 56) are clustered in the Asia clade (e.g., Thailand, Malaysia, Vietnam, Bangladesh, and Cambodia). Group III (ST 354) and V (ST 90) were found in a clade that included Asia, Australian, and Europe countries (e.g., China, Belgium, Madagascar, USA, and UK) (Figure. 4) . Myanmar’s three STs (ST 300; ST 56, and ST 416) were seen in same STs with Thailand STs, whereas the remaining STs had the distance in SLV, DLV, and TLV with Thailand STs. Myanmar common ST 90 exhibited the same STs as STs of USA, Australia, UK, and China and accounted for 60% (Figure. 4). There was a history in which Americans traveled, and Australian worked in World War II and in the construction of railway (namely Dead Railway construction from Burma to Thailand at the time of the Japanese colonization)

**Figure 4.**
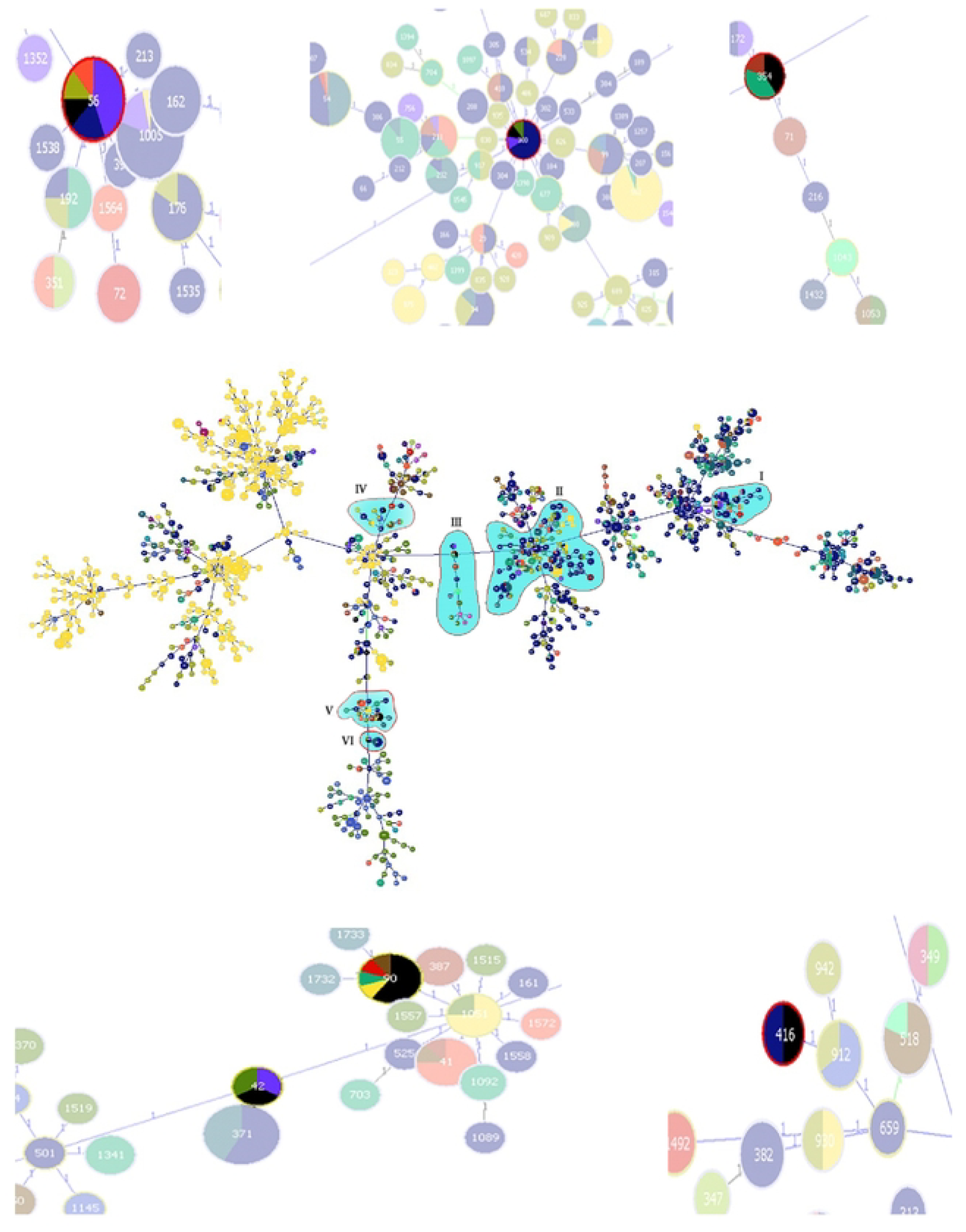
PHYLOViZ analysis of STs of Myanmar clinical and soil isolates showing the same cluster with global STs from PubMed datasets. **Group V and VI consisting of overlapped STs 90 (10 isolates) and overlapped STs 42 (3 isolates)**STs 90: (Black) Myanmar 60%(yellow). Australia 10%. (light Green), China 10%. (light red). UK 10%. (Brown), USA 10%. STs 42: (Green) India 33.33%, (Light blue) Bangladesh 33.33%. (Black) Myanmar 33.33%.**Group III consisting of overlapped STs 354 (5 isolates)** STs 354: (Black) Myanmar 40%. (light Green) China 40%. (Light Brown) Unknown 20%.Group 1 consisting of overlapped STs 56 (16 isolates) STs 56: (Light blue) Bangladesh 43.75% (Dark blue) Thailand 18.75% (Black) Myanmar 12.5%, (deep yellow) Cambodia 12.5%. (light red) Vietnam 12.5%.**Group II consisting of overlapped STs 300 (9 isolates)** STs 300: (Dark blue) Thailand 66,67%. (light blue) Bangladesh 11.11% (Black) Myanmar 11,11%. (Green) India 1 1.11%,**Group IV consisting of overlapped STs 416 (2 isolates)** STs 416: (Dark blue) Thailand 50% (Black) Myanmar 50%

## DISCUSSION

MLST is a flexible and powerful epidemiological tool to study the distribution and evolution of bacterial populations (12). It has been commonly used to characterize *B. pseudomallei* and analyze MLST data using phylogenetic software such as geoBURST and PHYLOViZ (27). It has been popular in the differentiation of genetic relatedness of bacteria globally. To our best knowledge, this study was the first research in the field of characterization of *B. pseudomallei* by using MLST from various geographical locations in Myanmar cost-effectively. This work was undertaken to know the STs of Myanmar and whether they correlated with other global STs of *B. pseudomallei*. It highlighted that Myanmar’s first published country (former name: Burma) had a migratory story.

In this study, 14 STs were identified, and the overall diversity of isolates was 0.6 STs/isolate. The high locus sequence diversity and nucleotide diversity (□) for STs showed high diversity among Myanmar isolates. When analyzed by the PHYLOViZ program, Myanmar STs were the same as the published STs by the group founder and were found as no clonal complex to each other among Myanmar STs. Topology and grouping of all Myanmar STs with other Asian, Europe, and Australian STs by MEGA exhibited that most Myanmar isolates grouped together in the Asian clade. Myanmar STs have not grouped units. Because there were a few STs identified in this study to be compared and determined with the grouping of STs globally on a large scale. ST 300 was already reported as a group founder in other studies. However, the STs of studied isolates from the present study were found as the same STs from the already published group founder.

Interestingly, Myanmar STs (ST 42; ST 416; ST 56; ST 354, and ST 300) identified in this study were found as the same historical STs published from MLST datasets. It was likely that Myanmar STs might be older and historical ones from Asian STs (28). It was assumed that Myanmar’s isolates might remain last more than 10 years ago, showing the historical colonization among global STs because it could not determine molecular characterization for *B. pseudomallei* due to a shortage of laboratory facilities and poor-resource settings at that time.

This study identified eight novel STs out of 21 STs, including five soil and sixteen clinical isolates (32). It was concerned with the ability to produce novel offspring genotypes. On the other hand, it may be caused by the action of genetic re-assortment of existing alleles (33, 34). The historical prevalence and dissemination of different *B. pseudomallei* genotypes might be reflected by the presence of different genotypes with various frequencies in the study area or due to the expansion of local STs that yielded new isolates with novel STs (28).

It was suggested that PHYLOViZ analysis revealed Myanmar isolates which showed more grouping with the Asian clade than the Australian clade. PHYLOViZ analysis further exhibited that within the Asian clade, Myanmar isolate groups were genetically close to isolates from China, India, Cambodia, Vietnam, Bangladesh, and Thailand.

It was found that many STs overlapped with these countries. Some overlapping STs (e.g., ST 42; ST 300; ST 56; ST 354) were first isolated in Asian countries, assuming the continued dissemination of bacteria among Myanmar and Asian countries. The predominate path and beginning of the Bay of Bengal cyclone are generally between India, Bangladesh, and Myanmar, directly through respective countries (36). One likely cause of strain similarity is tsunami attacks such as in Phangnga, a region in southern Thailand, causing melioidosis with the dispersal of *B. pseudomallei* and higher aerosol formation (37). Another hint for overlapping Myanmar STs with Thailand and Vietnam STs was likely that there was a history of human migration and population traveling (38).

Interestingly, ST 90 of *B. pseudomallei* were identified in both soil and human. Tontae Township, Yangon, was where ST 90 of soil isolates were isolated. There are a lot of industrial and economic zones in Yangon, describing densely populated districts. Interestingly, one ST 90 of a patient from Naypyitaw province (200 miles from Yangon) showed the same ST 90 of soil isolate from Yangon. It was hypothesized that it might be infected due to traveling history and probably due to inoculation or inhalation routes. The predominant pattern or sequence types might link the molecular association between soil and human isolates. Thus, the airborne transmission of melioidosis passes from the contaminated soil to aerosols after rainfall or storm and to humans in this endemic region (Yangon) (43).

Myanmar common ST 90 was already found as a sub-group of ST 1051 from most Australian STs. Interestingly, Myanmar common STs 90 were found as genetic relatedness to STs from the UK, Australia, USA, and China. To our knowledge, ST 90 was first isolated only from human resources in the USA in 2000. In this study, *B. pseudomallei* (ST 90) was later cultured from the soil and clinical cases in 2018. Surprisingly, there was no record of the year of isolation for Australia and China (39). It was likely that Myanmar seemed to be a reservoir to transmit *B. pseudomallei* locally because there was no report of which it was isolated from environmental sources in MLST datasets (40, 41). In the case of the correlation between Asia and Australian clades, it was questionable whether ST 90 was historical or not. From the history of Myanmar’s point of view, it was assumed that servicemen or allied prisoners came to Myanmar and worked together to build up a railway (namely the death railway from Myanmar to Thailand) under the colonization of Japan (23). It was noted that there were approximately 30,000 deaths in Than-Phyu-Zayat Township, Mon state, and Taukhyan Township, Yangon. Taken together, ST 90 was infected in the UK and USA, and patients with a travel history were described in the MLST database. (39). As soil ST90 was not yet isolated in those areas, it was not easy to assume whether it was correlated. For example, ST 60 showed the association between Thailand and Australia, but ST 60 of Australia was not isolated from the actual regions of Australia (42). Additional study is needed to determine the genetic relatedness between Myanmar and Australia ST (ST 90).

## Acknowledgments

We are grateful to all the laboratory practitioners in Myanmar who collected and stored the leftover samples. We express our gratitude to Dr. Chanwit Tribuddharat, Department of Microbiology, Faculty of Medicine Siriraj Hospital, Mahidol University for this publication. We are obligated to Dr. Narisara Chantratita from the Department of Microbiology and Immunology, Faculty of Tropical Medicine, Mahidol University.

